# DNA damage and macrophage infiltration in the ovaries of the long-lived GH deficient Ames Dwarf and the short-lived bGH transgenic mice

**DOI:** 10.1101/2020.01.19.911677

**Authors:** Tatiana Saccon, Monique Tomazele Rovani, Driele Neske Garcia, Jorgea Pradiee, Rafael Gianella Mondadori, Luis Augusto Xavier Cruz, Carlos Castilho Barros, Yimin Fang, Samuel McFadden, Andrzej Bartke, Michal M Masternak, Augusto Schneider

**Affiliations:** Centro de Desenvolvimento Tecnológico, Biotecnologia, Universidade Federal de Pelotas - RS, Brazil; Faculdade de Veterinária, Universidade Federal do Rio Grande do Sul, RS, Brazil; Faculdade de Veterinária, Universidade Federal de Pelotas, RS, Brazil; Instituto de Biologia, Universidade Federal de Pelotas, Pelotas, RS, Brazil; Faculdade de Nutrição, Universidade Federal de Pelotas, Pelotas, RS, Brazil; Departments of Internal Medicine, Southern Illinois University School of Medicine, Springfield, IL, USA; College of Medicine, Burnett School of Biomedical Sciences, University of Central Florida, Orlando, FL, USA

**Keywords:** Growth hormone, Insulin-like growth factor-1, DNA damage, macrophages, granulosa cells

## Abstract

**Objective:** The aim of the study was to evaluate the role of growth hormone (GH) in DNA damage, macrophage infiltration and the granulosa cells number of primordial and primary follicles.

**Methods:** For these experiments six groups of female mice were used. Four groups consisted of Ames dwarf (Prop-1^df^, df/df, n=12) and their normal littermates (N/df, n=12) mice, between sixteen and eighteen month-old, receiving GH (n=6 for df/df, and n=6 for N/df mice) or saline injections (n=6 for df/df, and n=6 for N/df mice). The other two groups consisted of ten to twelve-month-old bGH (n=6) and normal mice (N, n=6). Immunofluorescence for DNA damage (anti-γH2AX) and macrophage counting (anti-CD68) were performed. Granulosa cells of primordial and primary follicles were counted.

**Results:** Female df/df mice had lower γH2AX foci intensity in in both oocytes and granulosa cells of primordial and primary follicles (p<0.05), indicating less DNA double strand breaks (DSBs). In addition, GH treatment increased DSBs in both df/df and N/df mice. Inversely, bGH mice had higher quantity of DSBs in both oocytes and granulosa cells of primordial and primary follicles (p<0.05). Df/df mice showed ovarian tissue with less macrophage infiltration than N/df mice (p<0.05) and GH treatment increased macrophage infiltration (p<0.05). On the other hand, bGH mice had ovarian tissue with more macrophage infiltration compared to normal mice (p<0.05). Df/df mice had less granulosa cells on primordial and primary follicles than N/df mice (p<0.05). GH treatment did not affect the granulosa cells number (p>0.05). However, bGH mice had an increased number of granulosa cells on primordial and primary follicles compared to normal mice (p<0.05).

**Conclusion:** The current study points to the role of the GH/IGF-I axis in maintenance of oocyte DNA integrity and macrophage ovarian infiltration in mice.

## Introduction

Ames Dwarf mice (df/df) have a defective Prophet of Pit1 (Prop1) gene, which impairs anterior pituitary gland development, resulting in deficient growth hormone (GH) secretion [1]. As a result, df/df mice have very low levels of circulating insulin-like growth factor I (IGF-I), are significantly smaller and live around 30 to 65% longer than normal littermates (N/df) [2]. In contrast, transgenic mice overexpressing bovine GH (bGH) have elevated plasma levels of GH, resulting in increased circulating IGF-I levels and adult body size [3, 4]. The lifespan of bGH mice is approximately 50% shorter than for normal littermates [4]. Collectively, this evidence points to an important role of the GH/IGF-I axis in the aging process.

A progressive decline with aging and consequent depletion of the ovarian primordial follicle reserve is the main cause of the onset of menopause in women [5]. Along with the reduction in follicle numbers, the quality of the remaining oocytes enclosed in these follicles also decreases with age [5, 6]. It is well established that the age-related decline in fertility is paralleled by a decrease in the ovarian reserve and the increase in chromosomally abnormal conceptions [7]. One of the primary cause of the increased risk to miscarriages with age is frequently attributed to the exponential rise of the oocyte chromosome mis-segregations, leading to karyotypic imbalances and aneuploidies in the offspring [8]. Many of these karyotypic abnormalities result in spontaneous abortion in the first trimester and thus contribute to the high frequency of pregnancy loss during this time window [9]. However, advanced maternal age also poses an increased risk of serious complications that manifest in later pregnancy, these include miscarriage, late fetal and perinatal death, stillbirth, preterm and extreme preterm birth, low birth weight, placenta praevia and pre-eclampsia [10, 11].

The GH/IGF-I axis also has a central role in ovarian function [12–14]. Previous studies of our group have shown that middle-age and old GH-deficient df/df mice have a larger ovarian primordial follicle reserve than N/df mice [15, 16]. This indicates that increased longevity is correlated with increased reproductive lifespan. Furthermore, bGH mice had a smaller ovarian reserve than normal mice [17], showing that the GH/IGF-I axis has a central role in the reproductive lifespan. In spite of the evidences of the role of GH/IGF-I in the preservation of the ovarian reserve little is known about its effects on the other ovarian parameters as the mice ages.

Several factors can influence fertility during aging. The adverse effects of DNA damage caused by environmental factors on somatic cells have been extensively reported [18]. Mammalian oocytes enclosed in primordial follicles remain arrested in prophase I of meiosis up to several decades in some species, including humans [19, 20]. This long period of dormancy which oocyte are submit through increases the chances of accumulating DNA damage, as shown in female mice and human, both of which, accumulate double strand breaks (DSBs) in primordial follicle oocytes with aging [21]. Substantial damage occurring throughout meiosis can have serious consequences if an adequate cellular response is not activated, and may result in infertility or development of defective embryos that are unable to result in full-term pregnancy [22, 23]. As a reaction to DSBs, kinases are known to phosphorylate histone 2Ax (γH2AX) on serine 139 [24, 25]. This post-transcriptional modification at the lesion site provides a platform for the ataxia-telangiectasia mutated (ATM)–mediated DNA damage signaling pathway to regulates the repair of DNA DSBs [26].

In addition to DNA damage, inflammation also increases with age and can negatively affect fertility [27]. Interleukin 1 deficient female mice have more primordial follicles and increased fertility than control females [28]. One of our previous studies identified by RNASeq that old df/df mice have more than 150 down-regulated gene ontology terms related to the inflammatory/immune response compared to N/df mice [29], pointing to a role of inflammation in ovarian aging. It is well known that adipose tissue and blood levels of pro-inflammatory cytokines, interleukin 6 (IL-6) and tumor necrosis factor alpha (TNF-α) are reduced in df/df mice [30, 31]. This reduced inflammation is believed to be one of the main reasons for the increased longevity of df/df mice [32]. Given the background, the presence of inflammatory cells within the ovaries is an important parameter, as macrophages are the most numerous immune cells within the ovary [33]. Inflammation can result in oxidative stress, reducing cellular antioxidant capacity, leading to overproduction of free radicals that react with cell membrane fatty acids and proteins impairing their function permanently [34]. Furthermore, excess free radicals can increase DNA damage, a predisposing factor for several age-related disorders [35].

In mammalian ovaries, the majority of oocytes are meiotically arrested and surrounded by a layer of flattened somatic granulosa cells, a structure known as primordial follicle [36, 37]. For the purpose to ensure the proper reproductive lifespan, most of the oocytes are maintained in this quiescent state within primordial follicles [38]. Granulosa cells play a fundamental role in initiating the growth of primordial follicles [38]. The number of granulosa cells can be a useful marker of follicular activation and development [39]. Several molecular networks mediate the interaction between somatic cells and germ cells in controlling the development of dormant mammalian oocytes [38, 40]. For example, granulosa produced mammalian target of rapamycin (mTOR) stimulates Kit which in the oocyte stimulates the phosphoinositide 3-kinase (Pi3k)/protein kinase B (Akt1) pathway and the transcription factor Forkhead Box O3a (FoxO3a) phosphorylation resulting in primordial follicle activation [38, 41, 42]. We have previously shown that GH has a central role in primordial follicle activation through activation of FoxO3a [16], however, to date little is known about its effects on granulosa cells number in primordial and primary follicles.

Based on this evidence, the aim of the study was to evaluate the DNA damage, macrophage infiltration and the granulosa cells number of primordial and primary follicles in the ovaries of aged df/df and N/df mice treated with exogenous GH and bGH transgenic mice.

## Materials and methods

### Animals and treatment

For these experiments six groups of female mice were used. Four groups consisted of Ames dwarf (Prop-1^df^, df/df, n=12) and their normal littermates (N/df, n=12) mice, between sixteen and eighteen month-old, receiving GH (n=6 for df/df, and n=6 for N/df mice) or saline injections (n=6 for df/df, and n=6 for N/df mice). The other two groups consisted of ten to twelve-month-old bGH (n=6) and normal mice (N, n=6). Mice were maintained under temperature (22 ± 2°C) and humidity (40–60%) controlled conditions. All experiments were approved by the Ethics Committee for Animal Experimentation from the University of Southern Illinois, IL, USA. Mice treated with GH received recombinant porcine GH subcutaneous injections (4 µg/g of body weight; Alpharma, Inc., Victoria, Australia) twice daily, beginning at fourteen months of age for six weeks. Control mice received saline injections in the same way as GH treated mice. After six weeks of treatment, mice were kept two more weeks in the same conditions until euthanasia. The GH treatment used in the current study was proven effective for increasing serum IGF-I concentrations and body weight gain in previous studies using the same dose and treatment length in young (one month-old) [43, 44], middle age (five month-old) [45] and old mice (sixteen month-old)[43]. Body weights were measured before the first GH injection and at the end of the six-week treatment to confirm effectiveness of the treatment.

### Tissue collection and processing

The mice were anesthetized and euthanized after fasting for 12 h and the pair of ovaries was collected, dissected from surrounding adipose tissue and placed in 10% formalin buffered solution. After that, the ovaries were removed from the formalin solution, dehydrated in alcohol, cleared in xylene and embedded individually in Paraplast Plus (Sigma Chemical Company®, St. Louis, MO, USA). One ovary of the pair was then serially sectioned at 5µm using a semi-automated rotary microtome (RM2245, Leica Biosystems, San Diego, CA, USA). Sections were randomly selected for immunohistochemistry analysis using slides impregnated with 3% organosilane (Sigma Chemical Company®, St. Louis, MO, USA) in ethanol.

### Immunofluorescence

For immunofluorescence analysis, the ovarian samples were deparaffinized with xylene and rehydrated with graded alcohols. The primary monoclonal antibodies were obtained from Abcam (Abcam Plc, Cambrigde, UK) and diluted in 1.5% BSA solution. The anti-gamma H2AX (γH2AX) phospo S139 antibody (ab11174), to indicate DNA damage [21] and anti-CD68 antibody (ab955), to indicate the presence of macrophage [46] were used at a final dilution of 1:500. The blockage of the endogenous peroxidase activity was achieved with hydrogen peroxide blocking solution, while the antigen recovery was performed in humid heat, during 3 min after the boiling point, in citrate solution at pH 6.0. Non-specific background staining was reduced by covering the tissue sections that received protein block with BSA and goat serum. Thereafter, slides were incubated overnight with the primary antibody in a humid chamber at 4°C. The slides with anti-γH2AX and anti-CD68 antibodies were incubated for 1 hour with secondary Alexa Fluor® 488 (ab150113) and Hoechst (ab228550) for nuclei stanning during 15 min. The slides were mounted with a drop of mounting medium under coverslips. The images of the follicles were captured by confocal microscope (Olympus FluoView™ 1000). Fluorescence intensity quantification for γH2AX and macrophage counting was performed by image analysis software Image J® [47]. γH2AX and granulosa cells number was measured in 3 follicles/class/mouse. Macrophage counting was performed as a total number of macrophages per sections of ovary. Each mouse had a total of 4 random sections from the center of the ovary counted. The number of granulosa cells was counted as visible nuclei using the Hoechst (ab228550) for nuclei staining.

### Statistical analyzes

All statistical analyzes were performed using Graphpad Prism 6 (Graphpad Inc., La Jolla, CA, USA). Two-way ANOVA was used for comparing the fluorescence intensity, number of granulosa cells and macrophage number of df/df and N/df mice (effect of the genotype, treatment with GH and the interaction). T-test was performed for comparing fluorescence intensity, number of granulosa cells and macrophage number between bGH and normal mice. A P value lower than 0.05 was considered as statistically significant.

## Results

Female df/df mice had less DNA DSBs in primordial and primary follicles, as indicated by by lower fluorescence for γH2AX in oocytes from primordial (*p* <0.0001) and primary (*p* <0.0001) follicles compared to N/df mice. Also, df/df mice had reduced γH2AX intensity in granulosa cells of primordial (*p* = 0.0299) and primary (*p* = 0.0002) follicles. GH treatment increased DSBs in both df/df and N/df mice, oocytes from primordial (*p* <0.0001) and primary (*p* < 0.0001) follicles. Also, GH treatment increased DSBs in granulosa cells from primordial (*p* <0.0001) and primary (*p* <0.0001) follicles in both df/df and N/df mice. In the other hand, bGH mice had higher quantity of DSBs (Figure 1). The fluorescence intensity for γH2AX in oocytes from primordial (*p* = 0.0001) and primary (*p* <0.0001) follicles were higher in bGH mice compared to normal mice. Also, granulosa cells from primordial (*p* <0.0001) and primary (*p* = 0.0466) follicles had a higher fluorescence intensity for γH2AX in bGH mice (Figure 2). Representative images of anti-γH2AX intensity in oocyte and granulosa cells of primordial and primary follicles are shown in Figure 3.

**Figure 1.**
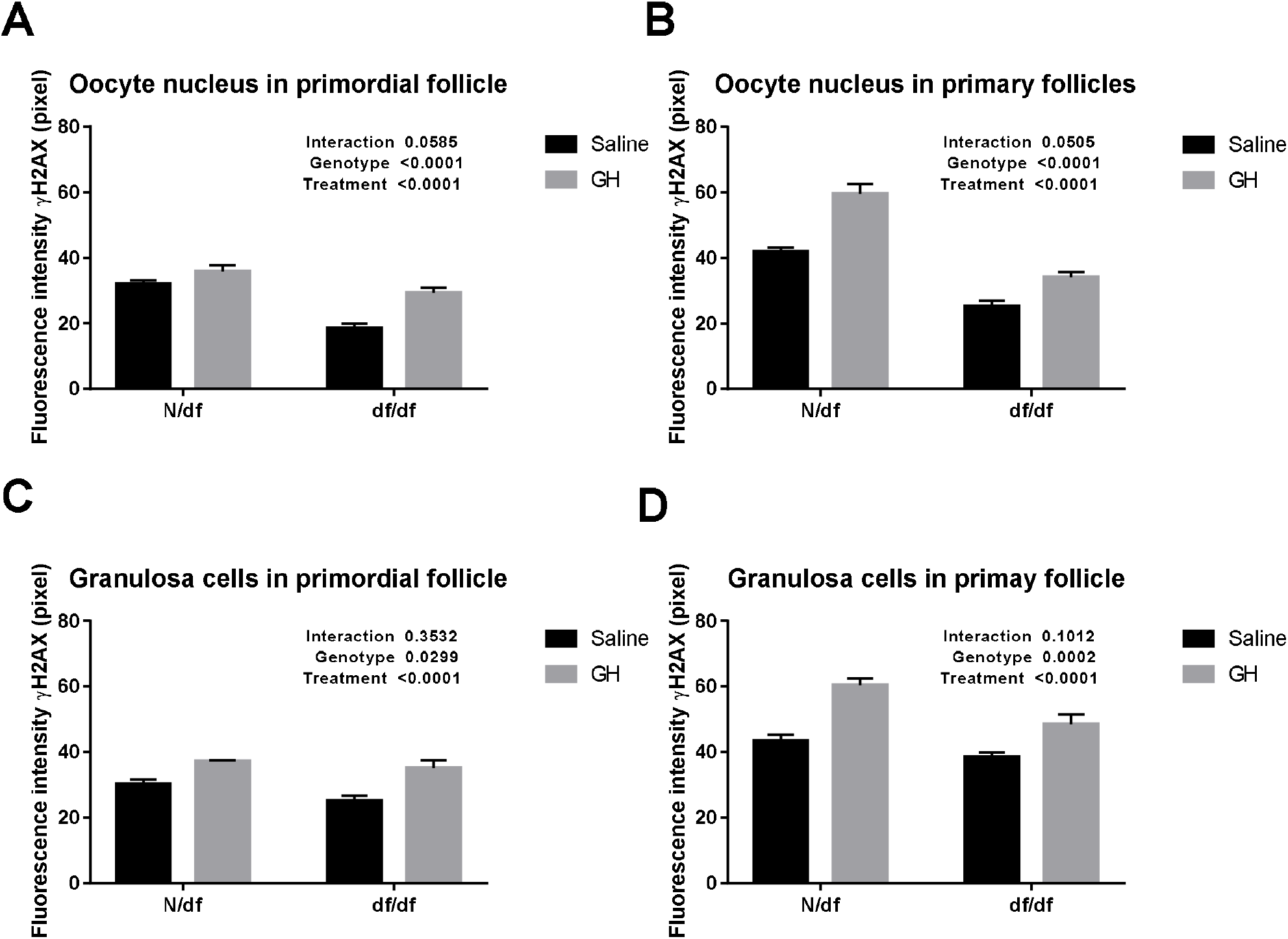
Fluorescence intensity of γH2AX immunostaining in oocytes nucleus of primordial follicle (n = 18, A), oocyte nucleus of primary follicle (n = 18, B) and granulosa cell nuclei from primordial (n = 18, C) and primary follicles (n = 18, D) of N/df and df/df mice treated with exogenous GH or saline. Data presented as media ± SEM.

**Figure 2.**
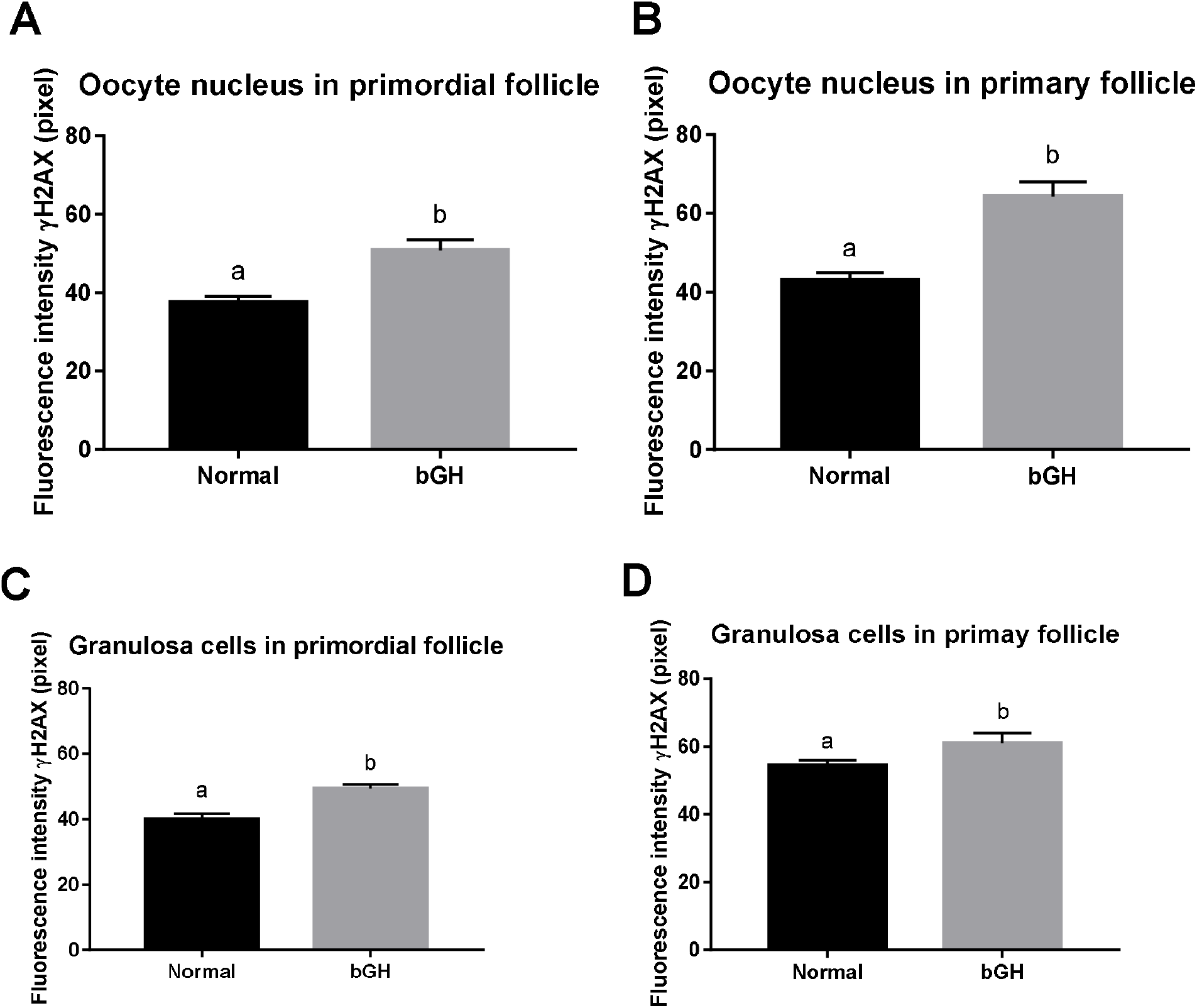
Fluorescence intensity of γH2AX immunostaining in oocytes nucleus of primordial follicle (n = 18, A), oocyte nucleus of primary follicle (n = 18, B) and granulosa cell nuclei from primordial (n = 18, C) and primary follicles (n = 18, D) of normal and bGH transgenic mice. Different letters indicate significant differences (*p* < 0.05). Data presented as media ± SEM.

**Figure 3.**
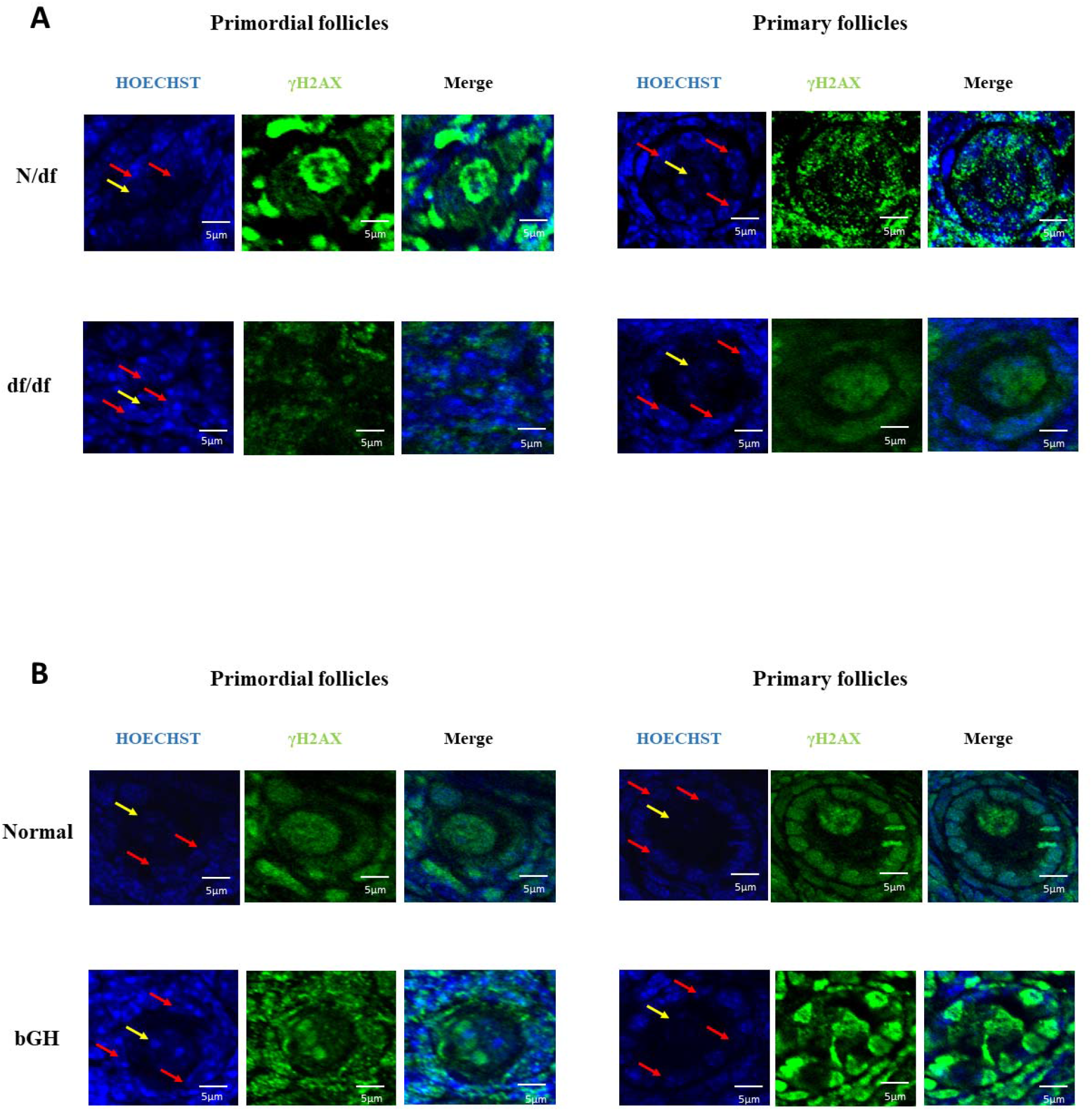
Representative images of immunofluorescence of anti-γH2AX in primordial and primary follicles of N/df and df/df mice (A); and normal and bGH mice (B). Blue images represent Hoechst (staining genetic material) and green represent anti-γH2AX staining γH2AX protein. Merged images are showed as both Hoechst and anti-γH2AX staining combination. Red arrows represent some granulosa cells and yellow arrow represents oocyte nucleus.

Df/df mice had ovarian tissue with the reduced macrophage infiltration (*p* <0.0001) and the GH treatment on df/df and N/df mice increased macrophage infiltration (*p* = 0.0007). On the other hand, bGH mice had ovarian tissue with increased macrophage infiltration (*p* <0.0001) (Figure 4).

**Figure 4.**
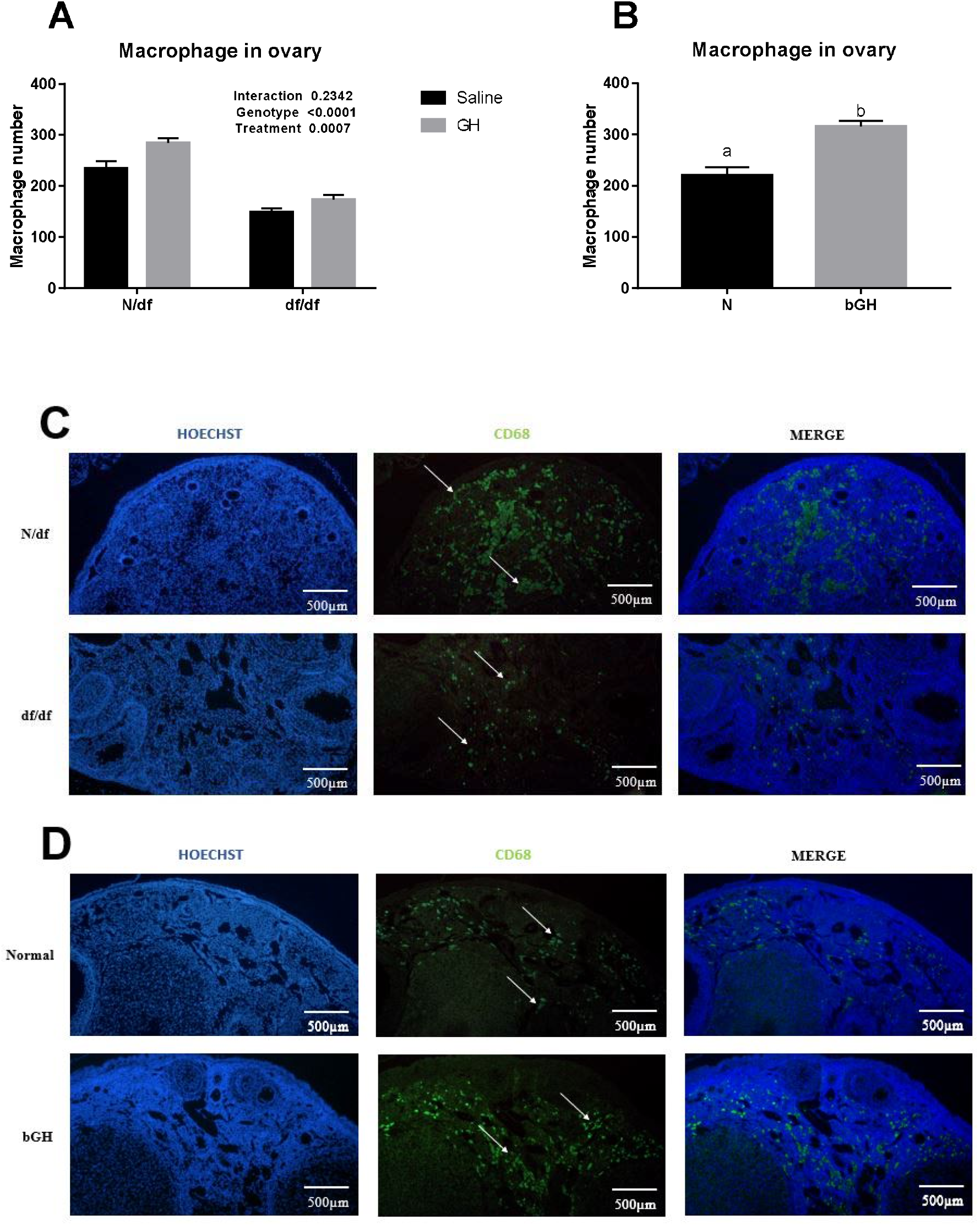
Macrophage number per ovarian section in N/df (n = 24) and df/df (n = 24) mice treated with exogenous GH or saline (A); and macrophage number on ovarian section of normal (n = 24) and bGH (n = 24) transgenic mice (B). Representative images of macrophage infiltration on ovarian section of N/df and df/df (C); normal and bGH transgenic mice (D). White arrow indicates macrophage staining. Different letters indicate significant differences (*p* < 0.05).

Df/df mice had less granulosa cells on primordial (*p* = 0.0002) and primary (*p* <0.0001) follicles than N/df mice. GH treatment did not change the number of granulosa cells of primordial and primary follicles (*p* >0.05). However, bGH mice had an increased number of granulosa cells on primordial (*p* <0.0001) and primary (*p* <0.0001) follicles compared to normal mice (Table 1).

**Table 1.** Number of granulosa cells of df/df and N/df mice treated with exogenous GH and bGH transgenic mice.

|  |  |  |  |  | *p*-Value<sup>a</sup> |  |  |
| --- | --- | --- | --- | --- | --- | --- | --- |
| Granulosa cell number (n=18) (SEM) |  |  |  |  | Genot. | Treat. | Genot*<br>Treat |
| Follicle Class | N/df |  | df/df |  |  |  |  |
|  | Saline | GH | Saline | GH |  |  |  |
| Primordial Follicles | 6.0 (±0.2) | 6.2(±0.2) | 4.8(±0.2) | 5.4(±0.2) | 0.0002* | 0.1389 | 0.5425 |
| Primary Follicles | 10.3 (±0.4) | 11.4(±0.6) | 8.8(±0.3) | 8.7(±0.3) | <0.0001* | 0.2796 | 0.1590 |
|  |  |  |  |  | *p*-Value<sup>b</sup> |  |  |
|  | Normal |  | bGH |  |  |  |  |
| Primordial Follicles | 5.4 (±0.4) |  | 9.9 (±1.1) |  | <0.0001* |  |  |
| Primary Follicles | 8.2 (±0.3) |  | 17.7 (±0.9) |  | <0.0001* |  |  |

<sup>a</sup>Two-way Anova test; <sup>b</sup>T-test; \*Significant P Value – p<0.05

## Discussion

The current study points to the role of the GH/IGF-I axis in maintenance of oocyte DNA integrity and ovarian macrophage infiltration. GH-deficient df/df mice are known for being smaller-sized and living longer and healthier life than normal littermate controls [2, 48], in the other hand, bGH mice are known for a bigger-sized body and short lifespan [4]. Our previous work [16], had shown that df/df mice have more ovarian primordial follicles than normal littermates, and that GH treatment increased primordial follicle activation and reduced the ovarian reserve. Additionally, we shown that bGH mice have decreased primordial follicle reserve. Therefore, our previous work shown increased ovarian reserve depletion promoted by GH, and our current findings also point to a role for GH in increasing oocyte and granulosa DNA DSBs and inflammation. This can suggest that beyond preserving the reserve for longer periods, GH/IGF-I deficiency also accelerates oocyte DNA damage which can impact fertility.

Oocyte and granulosa cells from primordial and primary follicles DNA DSBs were reduced in old GH-deficient df/df mice, while GH exogenous treatment increased accumulation of DNA DSBs. On the contrary, bGH mice had increased accumulation of DNA DSBs in oocytes and granulosa cells from primordial and primary follicles. DNA damage is a challenge that all somatic and germline cells are exposed during their lifetime [49]. Cells respond to DSBs as a serious threat to their integrity, activating DNA damage checkpoints (DDC) and reacting with a DNA damage response resulting in cell cycle arrest as a downstream effect, allowing activation of repair mechanisms [50]. DSBs trigger a DDC response by activating two major kinases, i.e. ATM and ataxia telangiectasia and Rad3-related (ATR) [51, 52]. Taking advantage of the cell cycle arrest, the cell can then repair the damaged DNA. At the DNA damage site, ATM gives rise to the phosphorylated form of the H2AX histone, which acts as a catalyst for the recruitment of the necessary checkpoint and repair factors [53]. Snell dwarf mice, endocrinologically identical to df/df mice, have increased cellular DNA repair capacity and upregulation of several DNA repair-related genes compared to normal littermates [54]. Our data, therefore, points to a role for the GH/IGF-I axis in DNA damage in mice oocytes and surrounding granulosa cells from the ovarian reserve. Mice and human oocytes accumulate DNA DSBs with age which contributes to reproductive aging [21]. Additionally, granulosa cells are essential for oocyte growth and activation, and others have also shown increased DNA DSBs in monkey granulosa cells with age [55]. Therefore, here for the first time we suggest that GH/IGF-I, which has a central role in somatic and gonadal aging, can also prevent accumulation of DNA damage, further contributing to the younger phenotype observed in GH-deficient mice. The decrease of DNA DSBs repair is also associated with accumulation of DSBs in human and mouse oocytes [21]. The expression of Breast Cancer 1 protein (BRCA1) and other key genes in the ATM-pathway decline with age in human oocytes, increasing DNA DSBs [21]. We previously observed a three-fold reduction in *BRCA1* gene expression with aging in N mice, although its expression did not change with aging in df/df mice [29], further contributing to DSB accumulation coordinated by GH/IGF-I axis. Overall, this work shows that GH/IGF-I deficiency beyond preserving the ovarian reserve for longer periods, can also contribute to maintaining oocyte DNA integrity, increasing the chances of successful pregnancy in older ages.

Old df/df mice exhibited reduced ovarian macrophage infiltration, and the treatment with GH in adult life increased the number of macrophages in the ovary. In the other hand, bGH mice had increased macrophage infiltration. These evidences suggest the GH/IGF-I axis as central to regulating ovarian macrophage infiltration. The long-living df/df, growth hormone receptor knockout mice (GHRKO) and calorie restricted mice have all been extensively characterized as having a reduced pro-inflammatory profile, which may represent one of the major mechanisms promoting increased insulin sensitivity and extended longevity in these mice [32]. Macrophages located in the ovaries, by secreting growth factors and/or cytokines, may play a synergistic role in stimulating cellular proliferation and follicle growth [56]. Some of the macrophage□derived factors that are known to impact follicular growth are hepatocyte growth factor (HGF), basic fibroblast growth factor (bFGF), tumor necrosis factor (TNF) α and β, and IGF-I [57–59]. On the other hand, prolonged exposure (both short-term and long-term) to a high-fat diet in young adult female mice reduced primordial follicle numbers, compromised fertility, produced higher systemic proinflammatory cytokine levels, and increased ovarian macrophage infiltration in the stroma, independent of obesity [46]. Pro-inflammatory Interleukin 1 deficient female mice have more primordial follicles and increased fertility than control females [28]. Also, LPS exposure in mice reduced the primordial follicle pool mediated by TLR4 [60], indicating the role of inflammation on ovarian reserve depletion. Our ovarian transcriptome study showed that the top 150 down-regulated terms between old df/df and old N mice were related to the inflammatory/immune response [29]. It was observed that macrophage chemotaxis, macrophage activation and macrophage differentiation were also among the top down-regulated biological processes in old df/df compared to old N mice. For that, these findings point to an association between the GH/IGF-1 axis and the inflammation on ovaries of old df/df mice. bGH mice have no data in ovarian inflammation, however, these transgenic mice have an increase pro-inflammatory profile in kidney [61] and age related increased pro-inflammatory markers in blood [62]. Another study with osteopetrotic mice showed reduced numbers of mature macrophages due to a natural mutation in the CSF□1 gene [63]. These osteopetrotic mice have reduced numbers of ovarian macrophages, which is not still clear if this is cause or effect of the decreased follicle growth on these mice. Our evidences provide an interesting connection between ovarian longevity and macrophage population coordinated by the GH/IGF-I axis, however, more studies are necessary to understand the role of macrophages in the ovarian aging.

The microenvironment within follicles and ovaries also could be influenced by GH/IGF-I deficiency. Particularly, granulosa cells and cumulus cells are intimately connected with oocytes, and both play critical roles in oocyte growth, follicular development and could also influence oocyte quality [64, 65]. Oocyte growth is accompanied by the active accumulation of mRNA, proteins, and lipids, and by modifications of chromatin configuration and DNA methylation status, and these molecular reactions require enough energy [66]. The number of granulosa cells in follicles is associated with the energy sufficiency of oocytes [66], which can affect oocyte and follicle growth. A complete set of molecular networks that mediate the interaction between somatic cells and germ cells in controlling the development of dormant mammalian oocytes had been described. The model proposed by Zhang et al. (2014) has two key steps. The first step is the mTOR complex 1 (mTORC1) signaling in granulosa cells that acts as the key decision-making process regarding whether or not a primordial follicle will be activated [38]. The second step involves the tightly regulated communication channel from the granulosa cells to the oocytes via KITL-KIT to trigger the awakening of the oocyte through FoxO3a phosphorylation [38]. It is proposed that these processes ensure the progressive activation of a limited number of primordial follicles throughout the reproductive lifespan. Thus, it is likely that follicular activation is initiated by molecular and cellular changes in the granulosa cells that are followed by awakening of the dormant oocytes. Our study shows that df/df mice had fewer granulosa cells surrounding oocytes than N/df mice in both primordial and primary follicles, supporting the late activation of primordial follicle observed for df/df mice in a previously study [16]. Also, bGH mice had a larger number of granulosa cells compared to normal mice, suggesting that granulosa cell number is an indication of successful follicle activation and development, since bGH mice presented less primordial follicles than normal mice [16]. This is in line with our previous work showing that FoxO3a activation is regulated by the GH/IGF-I axis in mice primordial and primary oocytes. Therefore, regulation of granulosa cell number by GH/IGF-I can be a critical point in determining ovarian lifespan.

## Conclusion

In conclusion, the present study indicates that GH/IGF-I is associated to oocyte and granulosa cell DNA damage and ovarian macrophage infiltration. Adding to our previous work, the current study demonstrates that beyond preserving the ovarian primordial reserve, GH/IGF-I deficiency prevents DNA damage and inflammation, both parameters linked to fertility in older ages. Overall, these important observations confirm, that delayed aging phenotype in these unique long-living mice, suggesting a beneficial characteristic of longevity in the reproductive organs of female df/df mice coordinated by GH/IGF-I levels.

## Acknowledgements

This is a preprint of an article published in Geroscience. The final authenticated version is available online at: https://doi.org/10.1007/s11357-021-00380-8.

